# Mosquito seasonality and trap type evaluation using routine surveillance data from New Orleans, Louisiana, United States

**DOI:** 10.1101/2025.09.25.678659

**Authors:** Nicole A. Scavo, Diana Reyes, Alexandros F. Pavlakis, Claudia Riegel, André B.B. Wilke

## Abstract

New Orleans, Louisiana, is a major gateway for the introduction of arboviruses into the United States due to high volumes of travel from arbovirus-endemic regions, the influx of migratory birds, and the presence of competent mosquito vectors. To respond to this increasing threat, the New Orleans Mosquito, Termite, and Rodent Control Board conducts routine and response-based mosquito surveillance. Three main types of traps are used in their surveillance system: gravid traps, CDC light traps, and BG-Sentinel traps. Understanding the capability of different trap types in assessing species richness, abundance, and community composition is instrumental in guiding surveillance efforts and responding to travel-associated and locally acquired arboviral infections effectively. This study aims to characterize the temporal dynamics of mosquito vector species in New Orleans, Louisiana, and to evaluate the effectiveness of BG-Sentinel, CDC light, and gravid traps in assessing species richness, abundance, and community composition. Alpha and beta diversity were compared based on trap type. Abundance, Richness, Shannon-Wiener diversity index, and evenness were compared using a Kruskal-Wallis Rank Sum test, followed by a post-hoc Dunn test with Bonferroni correction. Community composition was assessed using pairwise permutational multivariate analysis of variance (perMANOVA) and similarity percentage analysis (SIMPER), both with 10,000 permutations. Significant differences in mosquito abundance and diversity metrics were observed among trap types, indicating that trap choice strongly influences observed mosquito abundance, richness, community composition, and evenness. Gravid and CDC light traps, as well as BG-Sentinel and gravid traps, collected significantly different communities, driven mostly by higher *Culex quinquefasciatus* abundance in gravid traps. Differences between BG-Sentinel and CDC light traps were primarily driven by *Culex salinarius* and *Aedes vexans*, both more frequently collected by CDC light traps. Our results show that trap type significantly influences estimates of species abundance, richness, community composition, and evenness in New Orleans. These findings emphasize the importance of selecting appropriate trap types to generate accurate and actionable surveillance data, essential for guiding and optimizing mosquito control strategies aimed at preventing and responding to arbovirus outbreaks.

## Introduction

Mosquito-borne disease is a consistent threat to public health in the U.S. West Nile virus (WNV), transmitted by *Culex* mosquitoes, is the most prevalent mosquito-borne arbovirus in the United States. Current estimates indicate that approximately 7 million individuals have been infected by the WNV in the U.S. since its introduction in 1999 [1]. Local dengue transmission has also been reported in the United States [2]. Eighteen locally acquired dengue infections were reported in California in 2024: 14 in Los Angeles, 1 in San Bernardino, and 3 in San Diego [3]. Miami-Dade County, Florida, has reported 314 locally transmitted dengue virus infections from 2010 to 2024 [4]. Local dengue virus transmission has also been documented in Maricopa County, Arizona [5]. To monitor this increasing threat, mosquito surveillance programs are common throughout the U.S., with 734 mosquito control organizations across a variety of levels (e.g., state, county) [6]. These organizations rely on mosquito and arbovirus surveillance to inform mosquito control operations to prevent and respond to mosquito-borne disease outbreaks.

New Orleans, Louisiana, is a major metropolitan area in the U.S., with a population of approximately 370,000 spread over 900 km². It receives about 17 million tourists annually and serves as a hub for cruise ship traffic [7]. The high inflow of human movement to and from arbovirus-endemic regions positions New Orleans as an important gateway for the introduction of arboviruses into the U.S. The risk of local arbovirus transmission is amplified by the presence of competent mosquito vector species, although their distribution and abundance are heterogeneous depending on resource availability and environmental conditions. For instance, discarded tires are associated with increased abundances of *Aedes albopictus* and *Culex quinquefasciatus,* vectors of dengue and West Nile viruses, respectively [8,9]. Low-income neighborhoods often experience higher rates of tire dumping, increasing the availability of breeding sites for immature mosquitoes and intensifying the contact between mosquito vectors and human hosts in these areas [10].

To respond to this increasing threat, the New Orleans Mosquito, Termite, and Rodent Control Board conducts routine and response-based mosquito surveillance. Three main types of traps are used in their surveillance operation: gravid traps (John W. Hak Company, FL), Centers for Disease Control and Prevention miniature light traps (CDC light traps), and BG-Sentinel 2 traps (BioGents, Regensburg, Germany). Previous studies have shown that variations in trap attractiveness directly affect their ability to assess mosquito diversity and relative abundance reliably [11–14]. For instance, BG-Sentinel and CDC light traps are attractive to female mosquitoes that are host-seeking, whereas gravid traps are attractive to females that are seeking oviposition sites. Understanding the capability of different trap types in assessing species richness, abundance, and community composition is instrumental in guiding surveillance efforts and responding to travel-associated and locally acquired arboviral infections effectively. Mosquito intervention success depends on the timing of implementation and seasonality of the target species [15–17]. Therefore, the ability of the surveillance system to detect seasonal trends is essential for anticipating increases in mosquito abundance and optimizing the timing of intervention strategies [18].

We hypothesize that trap performance will differ by mosquito species due to species-specific behavioral, ecological, and physiological traits. This study aims to characterize the temporal dynamics of mosquito vector species in New Orleans, Louisiana, and to evaluate the effectiveness of BG-Sentinel, CDC light, and gravid traps in assessing species richness, abundance, and community composition.

## Materials & Methods

### Mosquito sampling

Mosquitoes used in the study were collected from January 2020 through December 2024 using CDC light traps baited with dry ice and gravid traps baited with hay and Ferti-lome Fish Emulsion Fertilizer (Bonham, TX). BG-Sentinel traps (Biogents, Regensburg, Germany) were deployed with BG-Lure in 2020; BG-Sentinel traps were not used in 2021. From 2022–2024, BG-Sentinel traps were baited with BG-Lure supplemented with CO₂ in the form of dry ice. The collected mosquitoes were identified to the species level using morphological characteristics and taxonomic keys [19–21]. Male mosquitoes were considered accidental catches and excluded from the analyses.

### Statistical Analysis

We set a threshold for trap nights to ensure consistent sampling effort and avoid inconsistencies in data collection. Traps with fewer than 10 trap nights per year (i.e., traps deployed fewer than 10 times for 24-hour periods during that year) were removed from the data set to maintain consistency with sampling effort. This filtered dataset was then used for all subsequent analyses. To ensure equal sampling effort for a comparison of alpha and beta diversity among trap types, a dataset was created in which at least 25 trap nights of each trap type (i.e., BG-Sentinel, CDC light, and gravid traps) per month were included in the analyses. Then, the data was randomly subsetted from the filtered dataset to have exactly 25 traps of each type per month to ensure equal trapping effort per trap type and to control for seasonality that also affects which species and how many individuals per species are collected throughout the year.

Alpha and beta diversity were compared based on trap type. Abundance (N), Richness (S), Shannon-Wiener diversity index (H’), and evenness (J) were compared using Kruskal-Wallis Rank Sum tests as the data was not normally distributed, and post-hoc Dunn tests with a Bonferroni correction for multiple tests used on the same data. Community composition was also compared among trap types using a pairwise permutational Multivariate Analysis of Variance (perMANOVA) with 10,000 permutations and Similarity Percentage Analysis (SIMPER) with 10,000 permutations.

## Results

A total of 1,020,749 female mosquitoes were collected across trap types, with 177,414 collected by BG Sentinel traps, 272,727 collected by CDC light traps, and 1,020,749 collected by gravid traps. Monthly trends of mosquitoes collected by gravid traps showed substantial heterogeneity. *Aedes aegypti* and *Ae. albopictus* exhibited well-defined seasonal patterns, with peaks in June and July, consistent across years despite varying relative abundance. *Culex quinquefasciatus* had the highest overall abundance, peaking from May to August, particularly in 2022 and 2024. *Culex coronator* had relatively low abundance peaking in June and July, except in 2024 when it peaked in May. *Culex nigripalpus* was more abundant in September and October 2024, while being low in other years. *Anopheles quadrimaculatus* peaked in May 2021, and *Anopheles crucians* had a moderate peak in abundance in May 2021, with relatively consistent low levels throughout other months and years (Figure 1).

**Figure 1.**
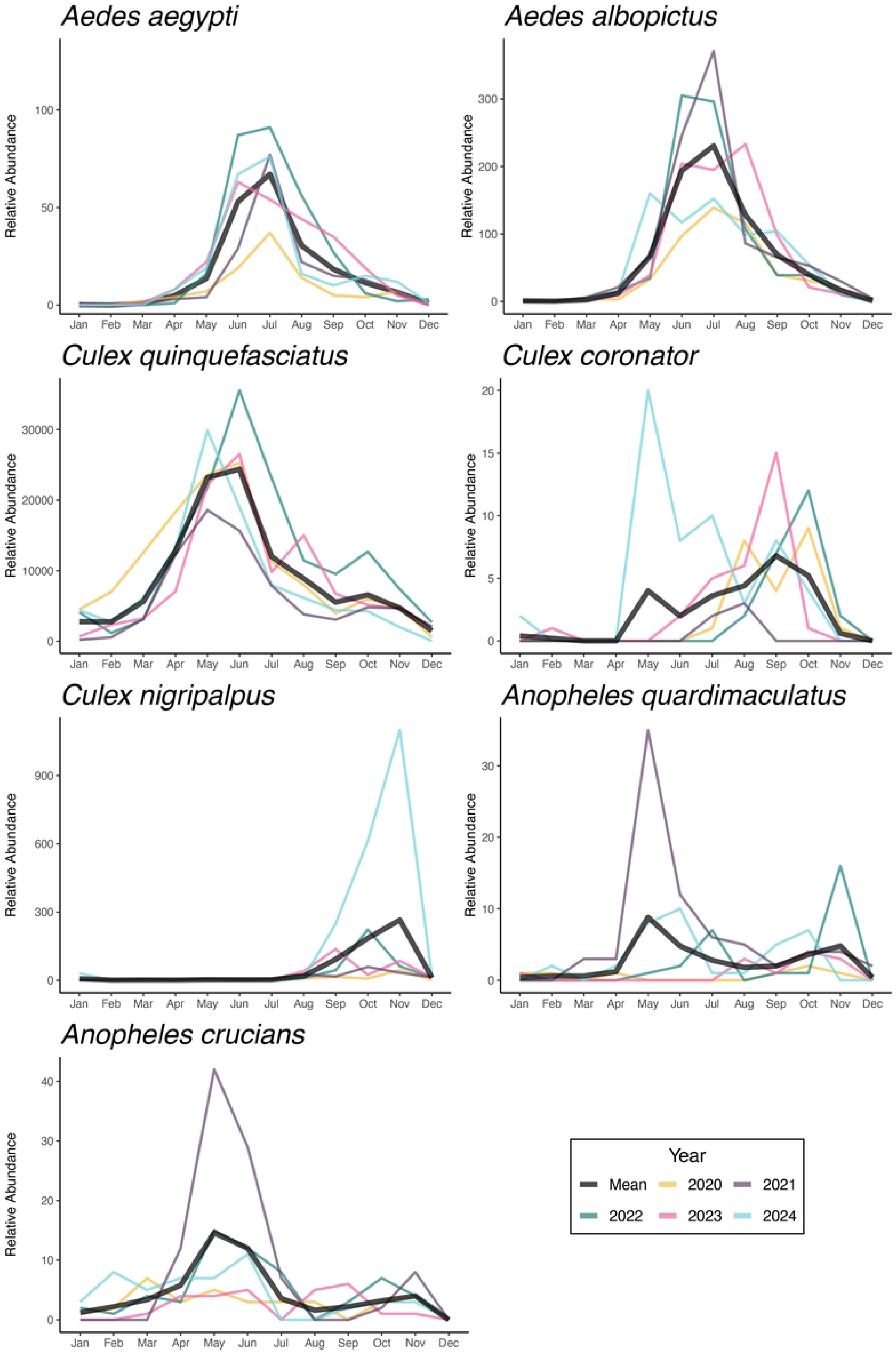
Population dynamics of mosquito vector species in New Orleans, Louisiana, collected using gravid traps. The relative abundance is shown year-round from 2020 to 2024. Individual years are shown in color, with the mean number of mosquitoes collected displayed in black.

The relative abundance of mosquitoes collected by the CDC light traps was also heterogeneous. *Aedes aegypti* exhibited seasonal peaks from May to August, with higher abundances in 2021. *Aedes albopictus* reached peak abundance in 2021, exhibiting two distinct peaks, one in June and another in October. *Culex quinquefasciatus* showed sustained activity throughout the year, with broad summer peaks most pronounced in 2020 and 2022. *Culex coronator* displayed an increase in abundance in August 2023, while remaining low in other years. *Culex nigripalpus* showed a sharp spike in relative abundance during August–September 2023, exceeding levels observed in different years. *Anopheles quadrimaculatus* peaked in May 2021, with lower and more stable abundance levels in subsequent years. *Anopheles crucians* showed high interannual variability, with elevated abundances from March to June in 2020 and 2022 (Figure 2).

**Figure 2.**
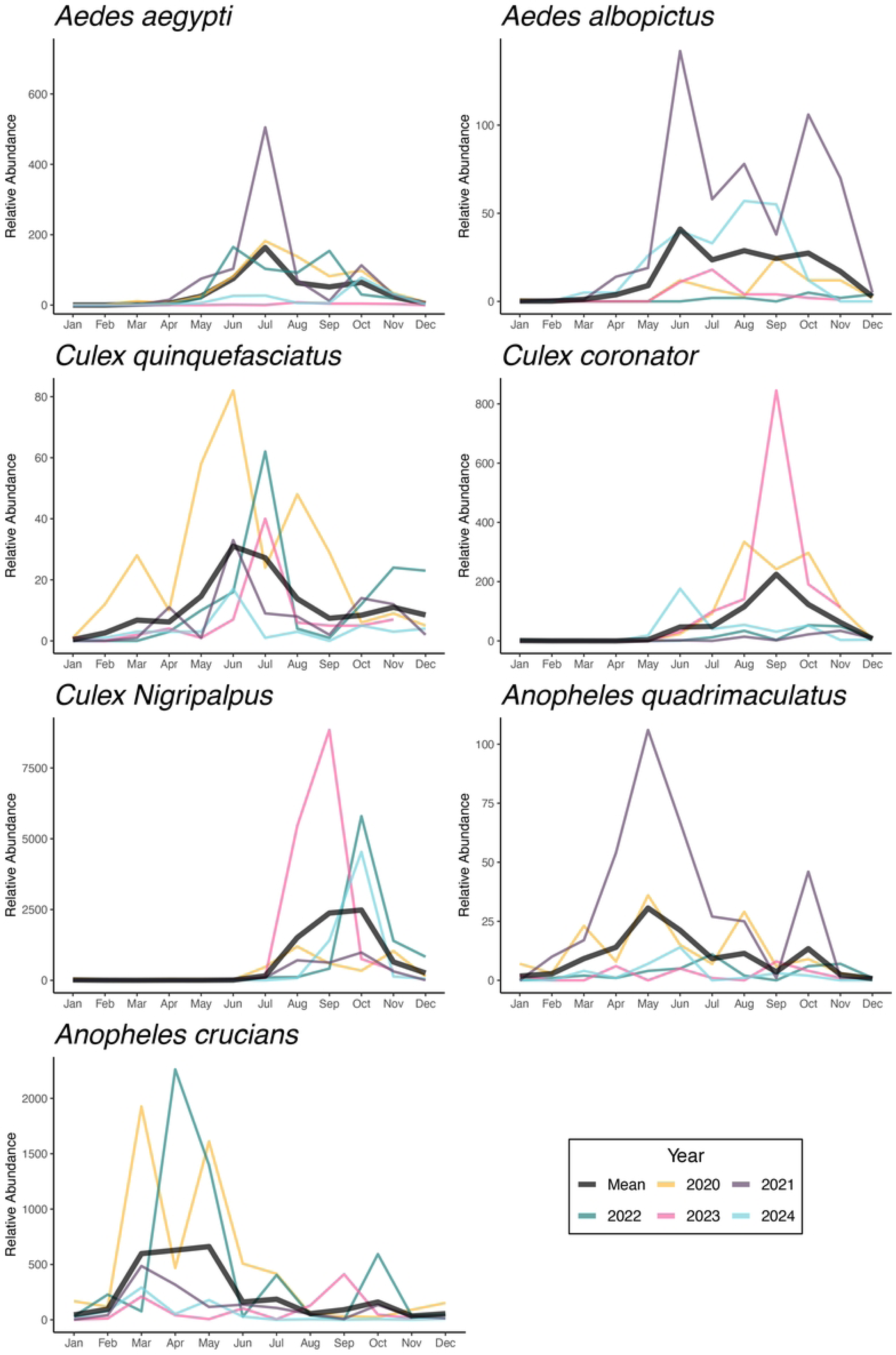
Population dynamics of mosquito vector species in New Orleans, Louisiana, collected using CDC light traps. The relative abundance is shown year-round from 2020 to 2024. Individual years are shown in color, with the mean number of mosquitoes collected displayed in black.

**Figure 3.**
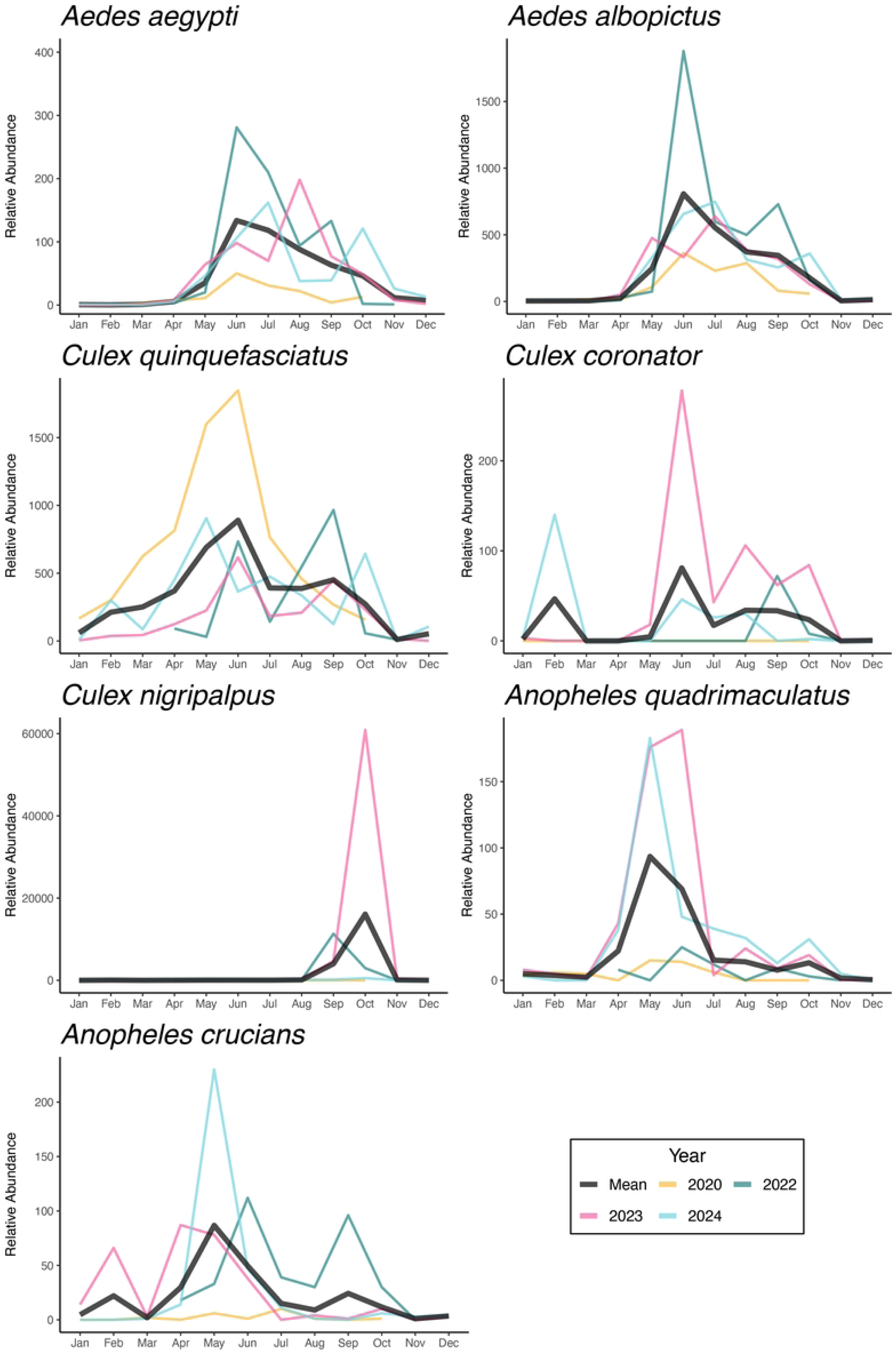
Population dynamics of mosquito vector species in New Orleans, Louisiana, collected using BG-Sentinel traps. The relative abundance is shown year-round from 2020 to 2024. Individual years are shown in color, with the mean number of mosquitoes collected displayed in black.

The relative abundance of mosquitoes collected using BG-Sentinel traps exhibited substantial temporal variability. *Aedes aegypti* and *Ae. albopictus* showed consistent seasonal peaks from May to July, with higher *Ae. albopictus* and *Ae. aegypti* abundance in 2022. *Culex quinquefasciatus* was more abundant from May to August, with the highest relative abundance in 2020 and consistent annual presence. *Culex coronator* had low abundance overall but showed sharp increases in 2023, particularly in June and July. *Culex nigripalpus* yielded a substantial abundance spike in September 2023, surpassing levels observed in all other years and species. *Anopheles quadrimaculatus* and *An. crucians* displayed seasonal increased activity from April to June, with *An. quadrimaculatus* peaking in 2023 and 2024, and *An. crucians* showing elevated abundances in 2024 and higher levels of variation in 2022.

Abundance and diversity measures showed significant differences among trap types. Relative mosquito abundance was significantly different among trap types (chi-sq = 203.9, df =2, p < 0.001). BG-Sentinel traps collected fewer mosquitoes than gravid and CDC light traps (BG-Sentinel and gravid: Z = −11.5, p < 0.001, BG-Sentinel and CDC light traps: Z = −13.0, p < 0.001, gravid and CDC light traps: Z = 1.5, p = 0.18) (Figure 4A). Species richness differed significantly among trap types (chi-sq = 957.0, df = 2, p < 0.001) with gravid traps yielding higher richness, followed by CDC and BG-Sentinel traps (BG-Sentinel and gravid traps: Z = −30.7, p < 0.001, BG-Sentinel and CDC light traps: Z = −12.0, p < 0.001, gravid and CDC light traps: Z = −18.7, p < 0.001) (Figure 4B). Evenness (chi-sq = 874.1, df =2, p < 0.001) and Shannon-Wiener diversity (chi-sq =447.5, df =2, p < 0.001) differed significantly among trap types, highest in CDC light traps, followed by BG-Sentinel traps and gravid traps (BG-Sentinel and gravid traps: Z = 26.6, p < 0.001, BG-Sentinel and CDC light traps: Z = 5.95, p < 0.001, gravid and CDC light traps: Z = - 22.9, p = 0.36) (Figure 4C). Shannon diversity was also significantly different among trap types (BG-Sentinel and gravid traps: Z =10.0, p < 0.001, BG-Sentinel and CDC light traps: Z = −11.1, p < 0.001, gravid and CDC light traps: Z = 21.1, p < 0.001), with the highest values for BG-Sentinel traps, followed by CDC light and gravid traps (Figure 4D).

**Figure 4.**
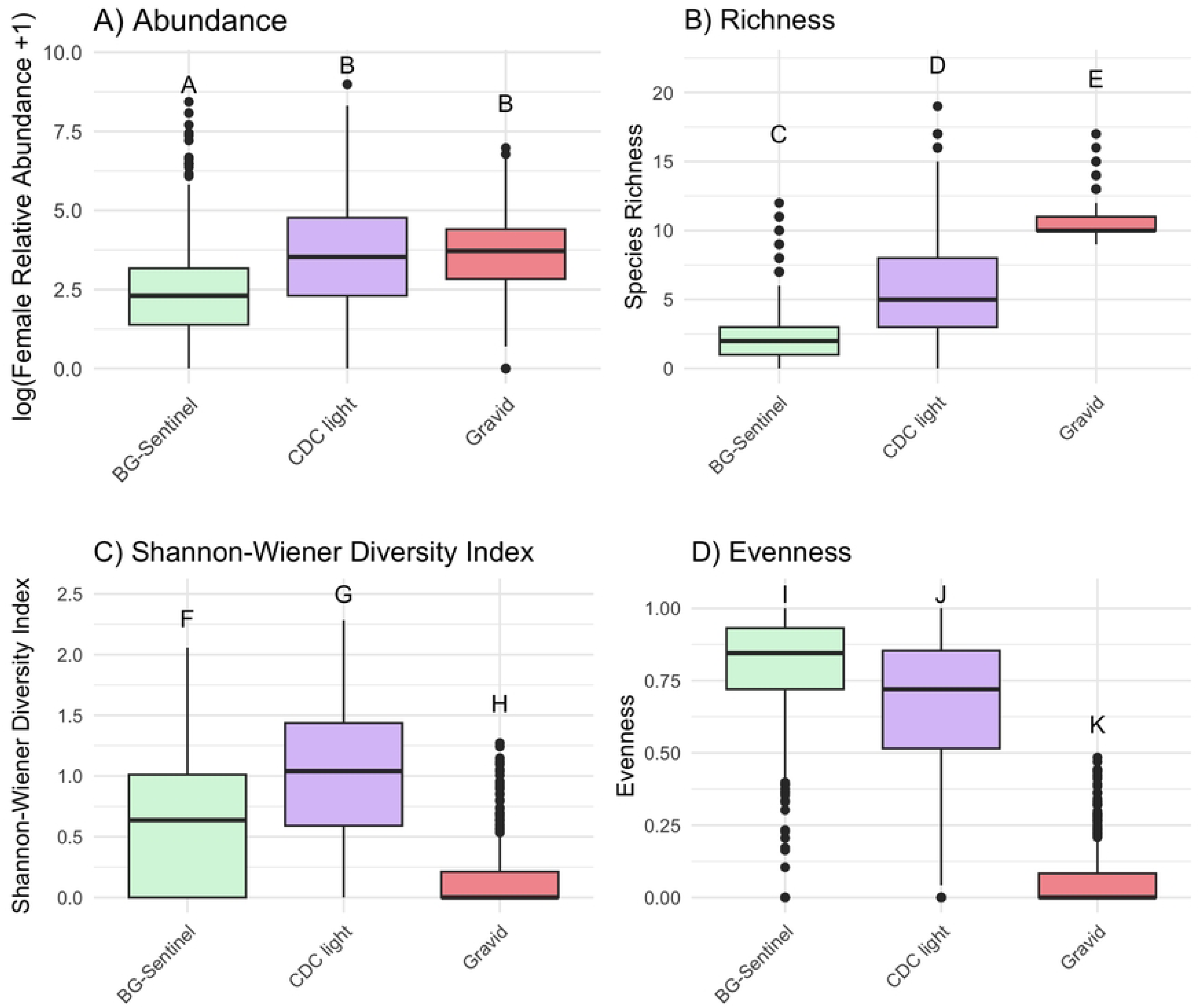
Mosquito abundance and diversity metrics compared across BG-Sentinel, CDC light, and gravid traps. (A) Abundance; (B) Species Richness; (C) Shannon-Wiener Diversity Index; and (D) Evenness. Letters indicate groups of significance (p < 0.05).

Community composition varied among the trap types. Gravid and CDC light traps collected significantly different communities (F = 354.1, p < 0.001). This difference was primarily driven by *Cx. quinquefasciatus,* which explained 45% of the difference between the two trap types and was more commonly collected by gravid traps (Table 1). BG-Sentinel and gravid traps also collected significantly different communities (F = 189.7, p < 0.001). Similarly, this difference was explained by differences in the abundance of *Cx. quinquefasciatus* collected by the traps (Table 2). BG-Sentinel and CDC light traps collected significantly different communities F = 116.13, p < 0.001). Most of the difference was explained by the abundances of *Cx. salinarius* and *Aedes vexans,* which were more commonly collected by CDC light traps (Table 3).

**Table 1.** Results from pairwise perMANOVA and SIMPER comparing community composition differences between CDC light and gravid traps. The perMANOVA and SIMPER were run with 10,000 permutations.

| **perMANOVA** | df | SumOfSqs | R<sup>2</sup> | F |  | *p* |  |
| --- | --- | --- | --- | --- | --- | --- | --- |
| model | 1 | 100.2 | 0.26 | 354.1 |  | **0.001** |  |
| residual | 1007 | 285 | 0.73 |  |  |  |  |
| **SIMPER** | average | sd | ratio | ava (CDC light) | avb (Gravid) | cumsum | *p* |
| *Culex quinquefasciatus* | 0.44 | 0.33 | 1.35 | 0.6 | 56.8 | 0.45 | **0.001** |
| *Culex salinarius* | 0.18 | 0.24 | 0.76 | 88.9 | 0.2 | 0.64 | **0.001** |
| *Aedes vexans* | 0.13 | 0.2 | 0.65 | 54.2 | 0 | 0.77 | **0.001** |
| *Culex nigripalpus* | 0.05 | 0.12 | 0.43 | 16 | 0.2 | 0.82 | **0.001** |
| *Anopheles crucians* | 0.02 | 0.07 | 0.43 | 9.5 | 0 | 0.85 | **0.001** |
| *Psorophora ferox* | 0.02 | 0.07 | 0.24 | 16.9 | 0 | 0.87 | **0.001** |

**Table 2.** Results from pairwise perMANOVA and SIMPER comparing community composition differences between BG Sentinel and CDC light traps. The perMANOVA and SIMPER were run with 10,000 permutations.

| **perMANOVA** | df | SumOfSqs | R<sup>2</sup> | F | *P* |  |  |
| --- | --- | --- | --- | --- | --- | --- | --- |
| Model | 1 | 41.54 | 0.1 | 116.13 | **0.001** |  |  |
| Residual | 1010 | 361.3 | 0.9 |  |  |  |  |
| **SIMPER Analysis** | average | sd | ratio | ava (BG-Sentinel) | avb (CDC light) | cumsum | *p* |
| *Culex salinarius* | 0.26 | 0.28 | 0.97 | 14.8 | 88.9 | 0.29 | **0.001** |
| *Aedes vexans* | 0.19 | 0.23 | 0.82 | 7.8 | 54.2 | 0.48 | **0.001** |
| *Culex quinquefasciatus* | 0.11 | 0.18 | 0.63 | 5.8 | 0.6 | 0.6 | 1 |
| *Culex nigripalpus* | 0.07 | 0.14 | 0.48 | 2 | 16 | 0.68 | **0.001** |
| *Aedes albopictus* | 0.05 | 0.12 | 0.49 | 3.4 | 0.8 | 0.75 | 1 |
| *Anopheles crucians* | 0.04 | 0.08 | 0.46 | 0.3 | 9.5 | 0.79 | **0.001** |

**Table 3.** Results from pairwise perMANOVA and SIMPER comparing community composition differences between BG Sentinel and gravid traps. The perMANOVA and SIMPER were run with 10,000 permutations.

| **perMANOVA** | df | SumOfSqs | R<sup>2</sup> | F | *p* |  |  |
| --- | --- | --- | --- | --- | --- | --- | --- |
| Model | 1 | 50.29 | 0.16 | 189.7 | **0.001** |  |  |
| residual | 1009 | 267.5 |  |  |  |  |  |
| **SIMPER Analysis** | average | sd | ratio | ava (BG-Sentinel) | avb (Gravid) | cumsum | *p* |
| *Culex quinquefasciatus* | 0.6 | 0.31 | 1.97 | 5.8 | 56.8 | 0.73 | **0.001** |
| *Culex salinarius* | 0.06 | 0.14 | 0.42 | 14.8 | 0.2 | 0.8 | 1 |
| *Aedes albopictus* | 0.05 | 0.1 | 0.52 | 3.4 | 0.4 | 0.87 | 1 |
| *Aedes vexans* | 0.05 | 0.12 | 0.39 | 7.8 | 0 | 0.92 | 1 |
| *Aedes aegypti* | 0.02 | 0.04 | 0.29 | 0.5 | 0.1 | 0.94 | 1 |
| *Culex nigripalpus* | 0.02 | 0.05 | 0.21 | 0.2 | 0.2 | 0.95 | 1 |

## Discussion

Global connectivity heightens the risk of arbovirus introduction into the U.S. [22]. To detect and respond to emerging arbovirus outbreaks, mosquito control districts in the U.S. rely on entomological surveillance [18]. However, mosquito species differ in their ecological niches, behavior, physiology, and habitat preferences, which directly influence their response to various trap types and attractants. Furthermore, mosquito traps are often designed to target specific species or life stages, introducing variation in collection efficiency and their ability to assess mosquito community composition.

Our results show that both abundance and diversity metrics differed significantly across trap types. Specifically, comparisons among BG-Sentinel, CDC light, and gravid traps yielded statistically significant differences in species richness, relative abundance, and community composition. Gravid and CDC light traps collected higher numbers of female mosquitoes than BG-Sentinel traps. Species richness was highest in gravid traps, followed by CDC light traps, with BG-Sentinel traps collecting the fewest species. The Shannon-Wiener diversity index was highest in CDC light traps, while gravid traps had the lowest diversity. Evenness was highest in BG-Sentinel traps, suggesting a more balanced distribution of species collected, whereas gravid traps were dominated by *Culex* species.

Results from the multivariate analysis showed that the mosquito community composition was significantly shaped by trap type. The SIMPER analysis indicated that gravid traps were dominated by oviposition-seeking *Cx. quinquefasciatus*, which alone accounted for most of the dissimilarity whenever gravid traps were compared, whereas host-seeking traps diverged mainly through *Cx. salinarius* and *Ae. vexans*, more abundant in CDC light traps, as well as contributions from *Ae. albopictus* in BG-Sentinel traps. These patterns confirm that gravid, BG-Sentinel, and CDC light traps collected partially overlapping but behavior-specific subsets of the mosquito assemblage; thus, combining at least one gravid and one host-seeking trap is recommended for balanced surveillance, while reliance on a single trap type risks inflating or underestimating the presence and abundance of key mosquito vector taxa. Although trap choice is important, more than 70 % of community variation remained unexplained, underscoring the need to incorporate environmental and spatial covariates for a deeper understanding of mosquito dynamics.

Mosquito assemblages collected by the three trap types revealed both overlapping and distinct patterns in mosquito population dynamics. Trap results were consistent in assessing seasonal peaks for *Ae. aegypti* and *Ae. albopictu*s, and the year-round relatively high abundance of *Cx. quinquefasciatus*. However, trap performance differed greatly for specific species. Gravid traps primarily collected *Culex* species, especially *Cx. quinquefasciatus*, with limited detection of *Aedes* and *Anopheles* mosquitoes. This aligns with their design, which targets gravid females seeking oviposition sites. CDC light traps, which use light and CO₂ as attractants, showed a more balanced collection across genera and revealed higher interannual variability, particularly for *Cx. coronator* and *Cx. nigripalpus*, suggesting greater sensitivity to fluctuations in environmental conditions. BG-Sentinel traps, optimized for host-seeking *Aedes* mosquitoes, collected the highest abundances of *Ae. aegypti* and *Ae. albopictus* and also detected peaks in *Cx. nigripalpus* in 2023, which were less evident in other trap types.

Our results show that, in New Orleans, *Ae. aegypti* peaked from May to August, while *Cx. quinquefasciatus* showed a broader season from April to September. Malaria vectors were detected throughout the year without a clear seasonal pattern, although *An. quadrimaculatus* peaked in April–July during 2023 and 2024. Regional comparisons highlight climate-driven differences: in arid Maricopa County, Arizona, *Ae. aegypti* peak activity is delayed (August–November) and *Cx. quinquefasciatus* had a bimodal abundance pattern, peaking in March–July and August– November [23]. In Los Angeles County, California, *Cx. quinquefasciatus* peaked in abundance in June–August and *Ae. aegypti* in July–November [24]. Subtropical Miami-Dade County, Florida, yielded similar patterns found in New Orleans for *Ae. aegypti* peaking from May to August, on the other hand, *Cx. quinquefasciatus* differed, peaking from November to May [25,26]. Malaria vectors showed weak seasonality with occasional increases in abundance [27]. Together, these findings indicate substantial heterogeneity, underscoring the need for locality-specific surveillance and intervention schedules taking into account local conditions and species assemblages.

BG-Sentinel trap deployment and baiting protocols in New Orleans varied across years. Changes in attractant regimes can alter trap efficiency and the taxonomic composition of collections, limiting comparisons. However, the comparison between BG-Sentinel and CDC light traps in New Orleans showed significantly different mosquito assemblages, with CDC light traps recording higher richness and BG-Sentinel traps greater evenness. On the other hand, a study in Miami-Dade County, Florida, detected no significant performance difference between BG-Sentinel and CDC light traps: each trap collected comparable numbers of mosquitoes across 23 species, with only one species unique to each trap [28]. Studies conducted in Germany and China found BG-Sentinel traps to be as effective as, or slightly more effective than, CDC light traps in assessing mosquito richness and abundance [14,29], whereas research in South Africa reported superior performance of CDC traps [30]. A study conducted in New Castle County, DE, and Salem County, NJ, found gravid traps to be more effective for West Nile virus surveillance than light traps or resting boxes [31]. In contrast, a study in Northeastern Florida found no statistically significant differences in mosquito assemblages between trap types [32]. These results underscore the importance of evaluating trap performance locally to inform surveillance design and refine mosquito control strategies, and strengthen arbovirus outbreak preparedness and response.

## Conclusion

Our results show significant differences in mosquito assemblages depending on trap type. These differences reflect the influence of mosquito behavior on trap performance. BG-Sentinel traps are more effective at estimating transmission risk, especially for anthropophilic species, while gravid traps are better suited for tracking *Culex* species, and CDC light traps for assessing overall diversity. Relying on a single trap type can bias species abundance estimates and underestimate key temporal or ecological patterns. These findings underscore the importance of selecting appropriate trap types when designing surveillance programs to ensure accurate and representative assessments of local mosquito populations.

## Acknowledgements

We thank the New Orleans Mosquito, Termite, and Rodent Control Board for conducting mosquito collection and species identification that supported this study.

